# Germline determinants of the somatic mutation landscape in 2,642 cancer genomes

**DOI:** 10.1101/208330

**Authors:** Sebastian M Waszak, Grace Tiao, Bin Zhu, Tobias Rausch, Francesc Muyas, Bernardo Rodríguez-Martín, Raquel Rabionet, Sergei Yakneen, Georgia Escaramis, Yilong Li, Natalie Saini, Steven A Roberts, German M Demidov, Esa Pitkänen, Olivier Delaneau, Jose Maria Heredia-Genestar, Joachim Weischenfeldt, Suyash S Shringarpure, Jieming Chen, Hidewaki Nakagawa, Ludmil B Alexandrov, Oliver Drechsel, L Jonathan Dursi, Ayellet V Segre, Erik Garrison, Serap Erkek, Nina Habermann, Lara Urban, Ekta Khurana, Andy Cafferkey, Shuto Hayashi, Seiya Imoto, Lauri A Aaltonen, Eva G Alvarez, Adrian Baez-Ortega, Matthew Bailey, Mattia Bosio, Alicia L Bruzos, Ivo Buchhalter, Carlos D. Bustamante, Claudia Calabrese, Anthony DiBiase, Mark Gerstein, Aliaksei Z Holik, Xing Hua, Kuan-lin Huang, Ivica Letunic, Leszek J Klimczak, Roelof Koster, Sushant Kumar, Mike McLellan, Jay Mashl, Lisa Mirabello, Steven Newhouse, Aparna Prasad, Gunnar Rätsch, Matthias Schlesner, Roland Schwarz, Pramod Sharma, Tal Shmaya, Nikos Sidiropoulos, Lei Song, Hana Susak, Tomas Tanskanen, Marta Tojo, David C Wedge, Mark Wright, Ying Wu, Kai Ye, Venkata D Yellapantula, Jorge Zamora, Atul J Butte, Gad Getz, Jared Simpson, Li Ding, Tomas Marques-Bonet, Arcadi Navarro, Alvis Brazma, Peter Campbell, Stephen J Chanock, Nilanjan Chatterjee, Oliver Stegle, Reiner Siebert, Stephan Ossowski, Olivier Harismendy, Dmitry A Gordenin, Jose MC Tubio, Francisco M De La Vega, Douglas F Easton, Xavier Estivill, Jan O Korbel, on behalf of the PCAWG Germline Working group^%^, and the ICGC/TCGA Pan-Cancer Analysis of Whole Genomes Network

## Abstract

Cancers develop through somatic mutagenesis, however germline genetic variation can markedly contribute to tumorigenesis via diverse mechanisms. We discovered and phased 88 million germline single nucleotide variants, short insertions/deletions, and large structural variants in whole genomes from 2,642 cancer patients, and employed this genomic resource to study genetic determinants of somatic mutagenesis across 39 cancer types. Our analyses implicate damaging germline variants in a variety of cancer predisposition and DNA damage response genes with specific somatic mutation patterns. Mutations in the *MBD4* DNA glycosylase gene showed association with elevated C>T mutagenesis at CpG dinucleotides, a ubiquitous mutational process acting across tissues. Analysis of somatic structural variation exposed complex rearrangement patterns, involving cycles of templated insertions and tandem duplications, in *BRCA1*-deficient tumours. Genome-wide association analysis implicated common genetic variation at the *APOBEC3* gene cluster with reduced basal levels of somatic mutagenesis attributable to APOBEC cytidine deaminases across cancer types. We further inferred over a hundred polymorphic L1/LINE elements with somatic retrotransposition activity in cancer. Our study highlights the major impact of rare and common germline variants on mutational landscapes in cancer.

## Introduction

Tumourigenesis involves somatic mutations arising as a result of exogenous and cell-intrinsic factors^1,2^. Recent efforts to sequence tumour genomes have revealed hundreds of somatically mutated genes in cancer^3-7^. They also uncovered a variety of mutational processes shaping the genomic landscapes of cancers^8-12^. These include G>T transversions attributed to tobacco carcinogens^13,14^, C>T and C>G substitutions attributed to APOBEC cytidine deaminase activity^15,16^ and spontaneous 5-methylcytosine deamination at CpG sites resulting in C>T transitions^17^. Mutational processes mediating genomic structural variants (SVs) are also active in cancer^18-23^, occasionally resulting in massively complex rearrangements (*e.g.* chromothripsis)^22,24-27^.

Genetic variants relevant to cancer additionally exist in the germline^28,29^, and many rare and common germline genetic variants have been implicated in cancer susceptibility^29,30^. Cancer predisposition genes frequently partake in fundamental cellular processes including cell cycle regulation and DNA repair^29^, the impairment of which may influence or augment the effects of somatic mutational processes^18,31^. Several studies have implicated rare and common germline cancer susceptibility variants in mutational processes^18,24,32-34^, patterns of selection of somatic variation^35,36^ and gene expression^37-40^. For example, carriers of pathogenic germline mutations in *BRCA1* and *BRCA2* typically exhibit characteristic somatic mutation patterns in breast, ovarian, pancreatic, and prostate tumours, and were found to exhibit increased rates of short insertions and deletions (indels), large deletions (in breast cancers with *BRCA2*-deficiency) and tandem duplications (in breast cancers with *BRCA1*-deficiency)^18,36,41-44^. Germline *TP53* mutations have been linked to chromothripsis in medulloblastoma^24^. Furthermore, evidence for germline variation influencing positive selection of somatic mutations has been revealed through analyses of single nucleotide polymorphism (SNP) arrays and whole exome sequencing (WES) data, exemplified by the identification of a haplotype associated with *PTEN* somatic mutation^35,36^. These findings emphasize the relevance of studying associations between germline and somatic variation, which could be markedly facilitated with the availability of unified germline and somatic variant calls in a substantial set of cancer genomes.

Here, we report on the efforts of the Pan-Cancer Analysis of Whole Genomes^45^ (PCAWG) germline variation working group, which constructed standardized whole genome sequencing (WGS)-based germline variant callsets in 2,642 cancer patients, to establish a reference resource of heritable genetic variants in donors from the International Cancer Genome Consortium (ICGC) and the Cancer

Genome Atlas (TCGA). We discovered 88 million germline variants in these donors, including SNVs, indels, and SVs. Integrative analyses of these germline variants with somatic SNVs, indels, and SVs seen in cancers obtained from these donors led us to uncover novel insights into the relationship of germline and somatic variation: We describe >100 active germline LINE/L1 elements mediating retrotransposition in cancer; uncover common variation modulating mutagenesis attributed to *APOBEC3B* across tumour types; and identify germline protein-truncating variants (PTV) in a variety of genes that associate with patterns of somatic mutation – including germline *MBD4* PTV variants, which our study implicates in a widespread mutational process.

## Results and Discussion

### A germline genome variation resource constructed from 2,642 cancer genomes

We employed several algorithms for germline variant detection in non-cancerous WGS samples from 2,642 PCAWG donors (**Fig. 1A** & **Methods**). The sequencing data, spanning 39 cancer types (**Table S1**) at a mean sequencing coverage of 39-fold, consists of WGS data from ICGC and TCGA studies^45^, which we analysed on high-performance and cloud computing platforms^46^. Integration of germline callsets by variant class yielded a site list with 80.1 million biallelic SNVs (harbouring two allelic states in the population), 5.9 million biallelic indels, 1.8 million multi-allelic short (<50bp-sized) variants, as well as SVs (defined as variants ≥50bp in size^47^) including 29,492 biallelic deletions and 23,855 mobile element insertions (MEIs) (**Table 1**). Our germline variant set incorporates genomic regions known to be difficult to sequence, including the HLA (human leukocyte antigen) loci that we analysed using ALPHLARD (**Supplementary Notes**). We further devised a methodology to statistically phase this germline variant set utilising 1000 Genomes Project^40,50^ haplotypes as a reference panel (**Methods**). The false discovery rate (FDR) of our site list was estimated to be <1% for SNVs, <1% for indels, <5% for deletions ≥50bp, <1% for MEIs, and 1% for multi-allelic short variants, respectively (**Table 1, Table S2**). Estimates for the non-reference genotype discordance were <1% and <3% for haplotype switch error rates at bi-allelic variants (excluding flip errors at rare reference panel sites), which we evaluated using haploid chromosomes from donor-matched tumour genomes (*i.e.*, somatic whole chromosome losses; see **Methods**).

**Table 1.** PCAWG germline variant callset based on ~39x (mean) coverage normal tissue whole genome sequencing read data from 2,642 cancer patients. FDR estimates are based on ultra-deep resequencing of randomly picked candidate variant sites in 48 PCAWG donors using custom sequence capture^#^ for SNPs and indels; intensity rank sum (IRS)^47^ testing on the basis of Genome-Wide Human SNP Array 6.0 datasets available for a subset (*N*=787) of PCAWG normal tissue samples* for SVs; and manual inspection of BAM files for a randomly selected subset of *Alu*, L1, SVA (for SINE-R–/VNTR–/*Alu*–based mobile elements), and endogenous retrovirus (ERV) insertions**. 798 variant sites within HLA regions (exons of *HLA-A, -B, -C, -DPA1, -DPB1, -DQA1, -DQB1, -DRB1*), median size of 1bp (median kbp per individual: 0.320) were separately genotyped using custom methodology (**Supplemental Notes**). *NA*, not assessed.

| Germline variant class | Number of sites | Median site size | Median kbp per individual | Site FDR |
| --- | --- | --- | --- | --- |
| Biallelic SNPs | 80,085,108 | 1bp | 3,536 | 0.0043<sup>#</sup> |
| Biallelic indels | 5,888,176 | 2bp | 789 | 0.0046<sup>#</sup> |
| Multiallelic short variants | 1,789,419 | 1bp | 689 | 0.0117<sup>#</sup> |
| Biallelic deletions ≥50bp | 29,492 | 2743bp | 2,805 | 0.0488* |
| *Alu* insertions | 19,641 | 312bp | 371 | 0.0071** |
| L1 insertions | 3,623 | 1704bp | 511 | 0.0102** |
| SVA insertions | 562 | 1275bp | 61 | 0.0172** |
| ERV insertions | 29 | *NA* | *NA* | 0.08** |
| Total | 87,816,050 |  |  |  |

**Figure 1.**
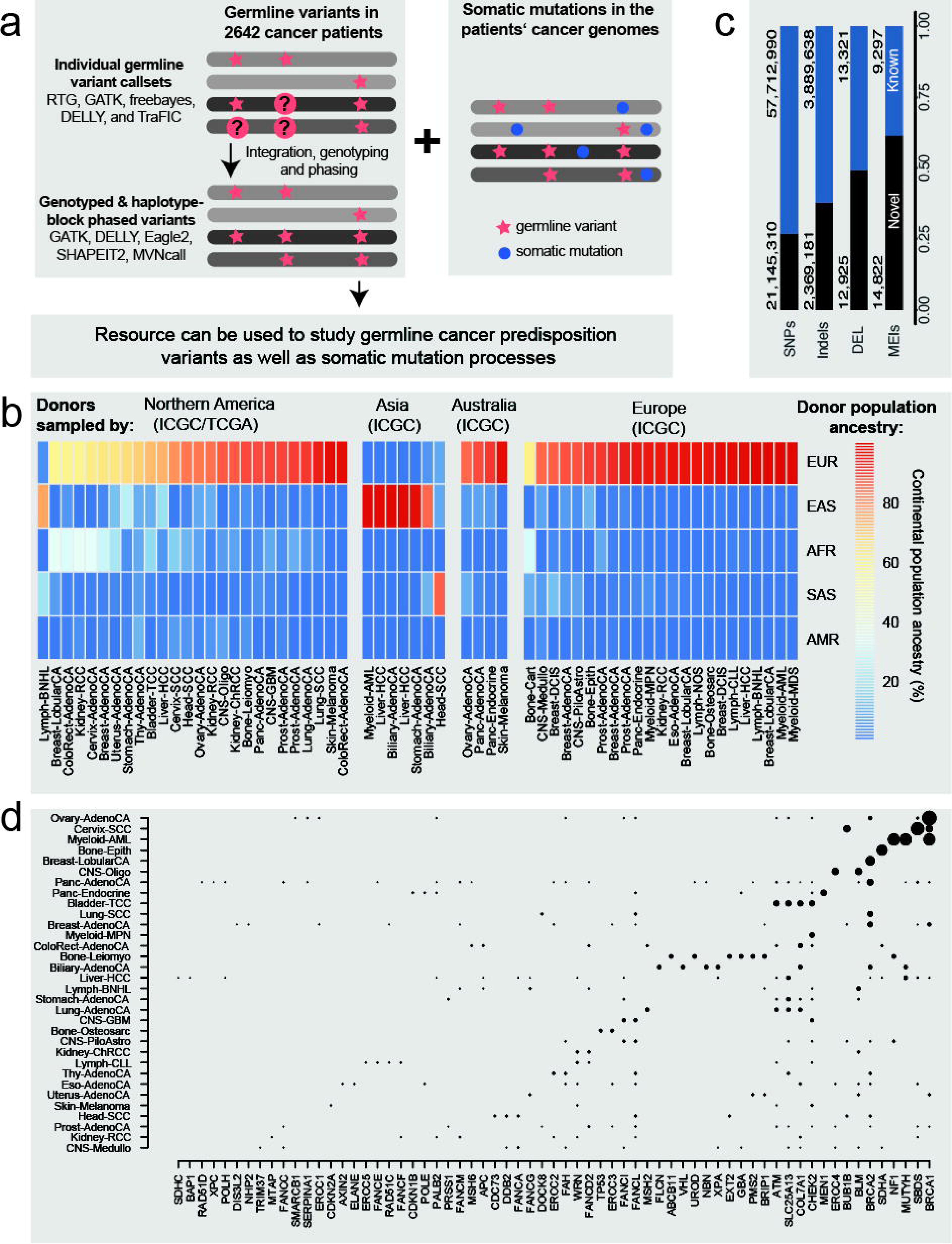
Approach taken and properties of the PCAWG germline variant callset. (**A**) Approach taken by the PCAWG germline variation working group: integration of germline and somatic variation called in 2,642 cancer patients. (**B**) Continental population ancestry of donors per tumour type (for tumour type abbreviations, see^45^) stratified by study origin (Northern America for TCGA studies; different continents for ICGC studies). AFR, African ancestry; AMR, native American ancestry; EAS, East Asian ancestry; EUR, European ancestry; SAS, South Asian ancestry. Data shown for all tumour types represented by *N*≥10 donors. (**C**) Novelty per germline variant class. Shown is the number of novel, autosomal variants compared to the union of variants from dbSNP, the 1000 Genomes Project and the Haplotype Reference Consortium. Multi-allelic variants were decomposed into individual variants and indels were left-aligned. DEL, deletions≥50bp. (**D**) Germline loss-of-function (PTV) variants in cancer predispoisiton genes, by histology (blob diameters scale with the number of carriers per tumor type, from 1 to 14).

We inferred donor population ancestry using a supervised version of the ADMIXTURE algorithm (**Methods**). The majority of PCAWG donors are of European (~75%) and East Asian (~15%) ancestry, reflecting the geographical distribution of ICGC and TCGA projects^45^ (**Figure 1B, Extended Data Fig. 1**). Most germline variant sites (81.4%) represented rare alleles with minor allele frequency (MAF) <1%, and 23.5M (27.6%) of these were novel with respect to widely used DNA sequence variation archives (**Figure 1C, Extended Data Fig. 2**). And while only 35,459 (0.2%) of all common autosomal SNVs/indels (MAF≥1%) were novel with respect to sequence variation archives, our data notably comprise >250,000 common alleles not previously incorporated into public haplotype reference datasets, which in the future can be utilised in genome-wide association studies (GWAS) focussing on cancer and other phenotypes via imputation^40^ (**Extended Data Fig. 2**).

**Figure 2.**
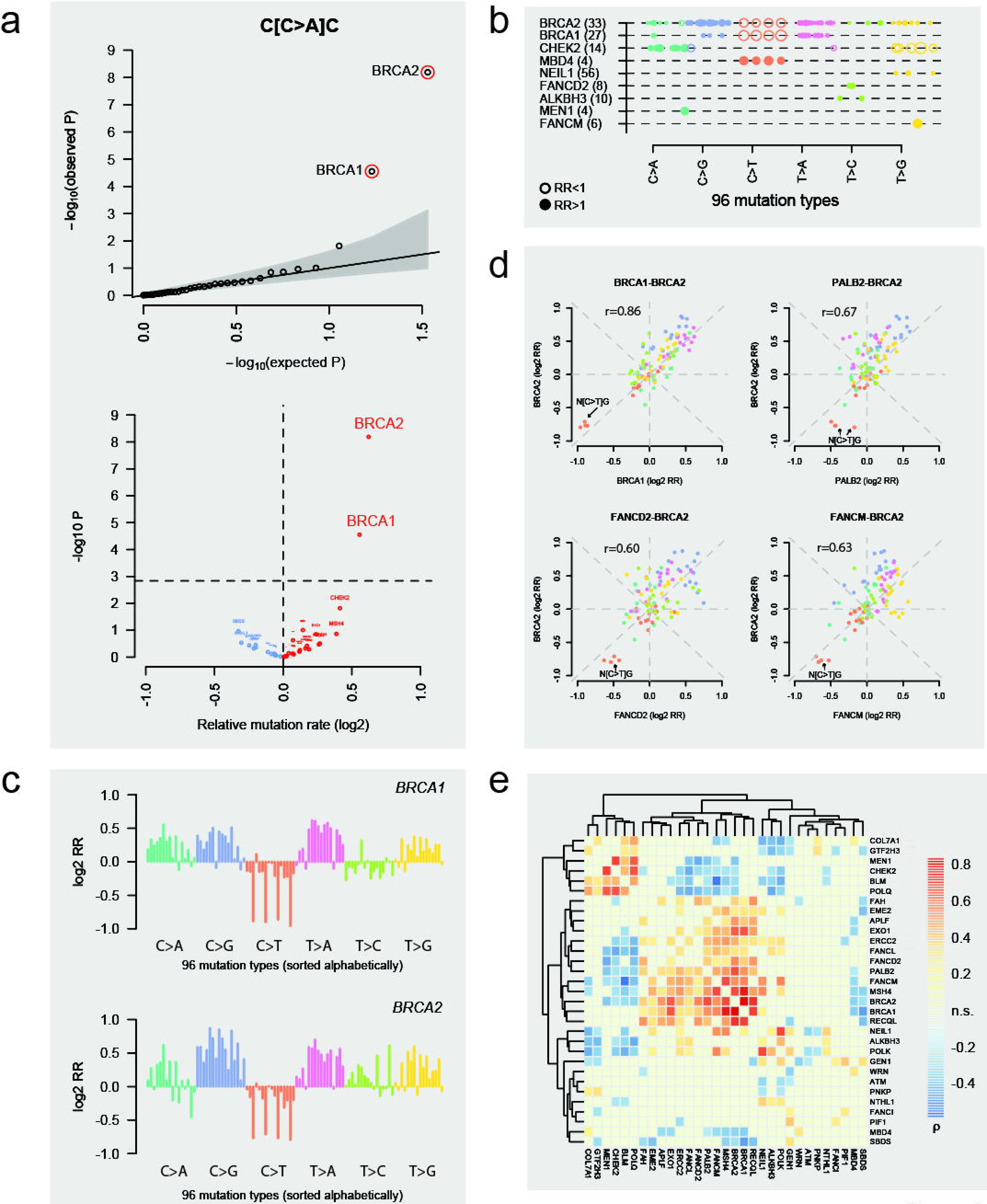
Association of germline gene loss-of-function (PTV) variants with somatic mutation types. **(A)** Quantile-quantile (QQ) plot and volcano plot, exemplified for a single mutation type (C[C>A]C). (**B**) Summary of mutation types seen at 10% FDR (computed over all candidate genes represented by *N*≥4 germline PTV carriers), where 96 mutation types are sorted alphabetically. Closed circles: relative enrichment; open circles: relative depletion. Numbers in parentheses show the number of germline PTV carriers found by candidate gene. Blob diameters scale with log_2_ relative rates (RR) for individual mutation types, scales shown range from 0.5 to 1.2. Closed circles reflect positive, and open circles negative values. (**C**) PTV-specific mutation type spectra (briefly, mutation spectra) of *BRCA1* and *BRCA2*. Relative rates (RR) shown are from our additive model, and control for potential confounders. (**D**) Pair-wise correlation shown for several selected mutation spectra of genes involved in HR and ICL repair. (**E**) Hierarchical clustering of mutation spectra, shown for all candidate genes with at least *N*=4 germline PTV carriers.

We identified hundreds of likely deleterious frameshift, nonsense, and splice site disrupting variants leading to protein truncation (PTV) of 109 known autosomal cancer predisposition genes^29^ (**Methods**), affecting 11% of PCAWG donors (**Figure 1D & Extended Data Fig. 3, Table S3)**. Germline PTVs among 183 DNA damage response genes^48^ without a presently established link to cancer risk were seen in another 20% of donors (**Table S3**). Testing for enrichment of rare variants with inferred functional consequence^49^ amongst annotated non-coding regulatory regions for this collective set of 292 cancer predisposition and DNA damage response genes did not reveal evidence for enrichment (**Supplementary Notes**).

**Figure 3.**
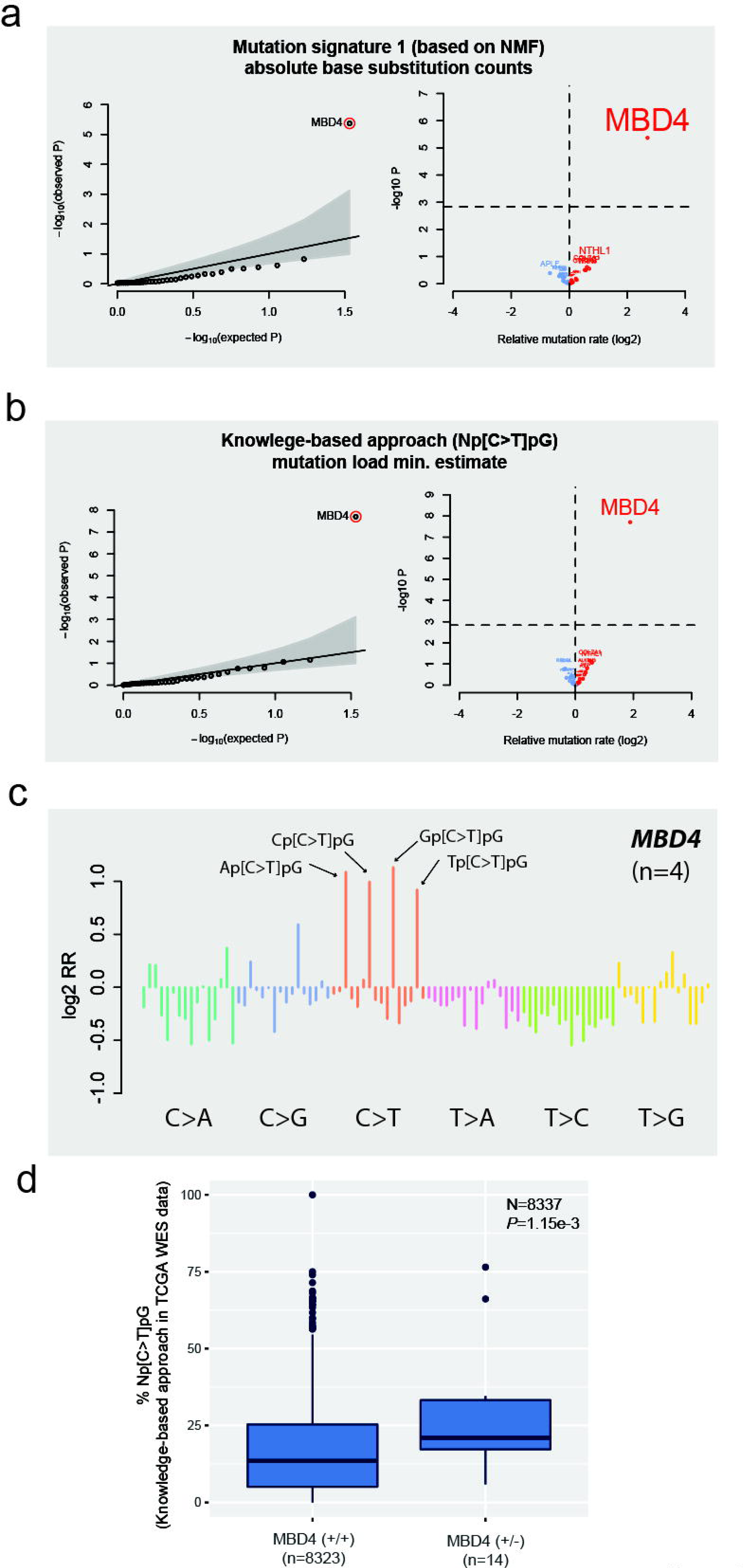
Association of germline PTVs in *MBD4* with elevated rates of somatic C>T mutation at CpG dinucleotides. **(A)** QQ and volcano plot showing association of *MBD4* with mutation signature 1 estimates across PCAWG histologies (European donors). **(B)** QQ and volcano plot showing association of *MBD4* with C>T mutations at CpG dinucleotides inferred using a knowledge-based approach (by searching for C>T mutations with NpCpGs, in European donors). **(C)** Visualization of the somatic mutation spectrum observed in *MBD4* germline PTV carriers. **(D)** Replication of *MBD4* association signal using TCGA exome sequencing data. A box plot depicts percentages of CpG>TpG mutations in tumours developing in *MBD4* germline heterozygous PTV carriers versus non-carriers.

As may be expected, previously reported cancer susceptibility SNPs identified though GWAS^50-57^ exhibited enrichment for tumour types in which those SNPs were originally identified. This included prostate cancer, chronic lymphocytic leukaemia (CLL), and melanoma, where polygenic risk scores (PRSs) showed elevation within the respective disease entity (*P*<0.001, Benjamini Hochberg-corrected Wilcoxon test; **Extended Data Fig. 3 & Supplementary Notes**). Furthermore, patients with oesophageal adenocarcinoma showed PRSs indicative of susceptibility to melanoma suggesting potential commonalities in disease pathways. In CLL patients, high CLL-specific PRSs were correlated with young age at diagnosis (r=-0.23, *P*=0.02, Spearman rank correlation; **Extended Data Fig. 3**).

### Germline PTV association with mutation types implicates several genes in mutagenesis

We next focused on investigating genetic determinants for somatic mutagenesis^45^. We performed rare variant burden tests (*i.e.* rare variant association analysis) to investigate the relationship of gene PTVs and somatic mutagenesis across cancers represented in PCAWG. To limit the number of hypotheses tested, we restricted our investigation to germline PTVs inferred amongst the 292 candidate genes (**Table S3**), with the reasoning that previous studies have successfully linked cancer predisposition genes and DNA damage response genes with patterns of somatic DNA alteration^24,32^. Unless stated otherwise, analyses presented in the following were confined to donors of European ancestry and accounted for additional potential demographic, histological, and technical confounders (**Methods**).

We first investigated single nucleotide substitutions, which we stratified by their local sequence context to account for the known sequence specificity of mutagenic processes^16,18,44^. This was achieved by considering in addition to each base substation, nucleotides at 5’ and 3’, which yielded trinucleotides corresponding to 96 mutation types^16,44^ (**Fig. 2A**). We identified nine genes displaying association between rare germline variants and the rate of substitution of at least one mutation type (**Fig. 2AB**), when adjusting FDR to 10% using the Benjamini-Hochberg procedure. These included genes with roles in (or interacting with proteins involved in) homologous recombination (HR) repair and DNA interstrand crosslink (ICL) repair (*BRCA1*, *BRCA2*, *FANCD2*, *FANCM*), base excision repair (*NEIL1* and *MBD4*), and other DNA damage-related cellular processes (*MEN1, CHEK2, ALKBH3*). Each of these genes showed enrichment or depletion of specific mutation types in conjunction with PTVs. For example, *CHEK2* germline PTVs were associated with increased C>A and decreased T>G substitution rates, whereas *BRCA2*, *NEIL1* and *FANCM* germline PTV carriers showed increased T>G substitution rates (**Fig. 2BC**).

For some genes, the relative rates of somatic mutation type enrichment and depletion seen in germline PTV carriers, which can be summarized in the form of somatic mutation type spectra (briefly: mutation spectra; **Fig. 2C**), appeared to be broadly similar. For example, the mutational spectra of tumours arising in *BRCA1*, *FANCD2*, and *FANCM* germline PTV carriers were highly correlated (Spearman’s *r*≥0.6; **Fig. 2D**) to tumours arising in *BRCA2* germline PTV carriers. To more closely investigate similarities and differences in mutation spectra we performed hierarchical clustering of genes in our list of 292 candidate loci for which at least four European germline PTV carriers existed in PCAWG. This analysis lends further support to the similarity in mutation spectra of these four genes and let us to infer additional genes, including *PALB2* and *FANCL* that have known roles in HR and ICL repair pathways^58,59^, with correlated mutation spectra (**Fig. 2DE**)

Defects in *BRCA1* and *BRCA2* have previously been implicated in an overrepresentation of mutations attributed to COSMIC mutation signature 3 (presumed to be mediated by a failure of DNA double-strand break-repair by HR)^44^, prompting us to relate germline PTVs in these genes with NMF-based signature 3 estimates^45^. We detected signature 3 in 84% of *BRCA1* and 73% of *BRCA2* germline PTV variant carriers in PCAWG. Overall, 92% (46/50) of tumours with biallelic inactivation of *BRCA1*/*BRCA2* (**Table S4, Extended Data Fig. 4**) were positive for signature 3 mutations and included ovarian (n=16), pancreatic (13), breast (12), prostate (2), and endometrial (1) cancers. Interestingly, none (0/8) of the monoallelic *BRCA1/BRCA2* cases exhibited signature 3 mutations (P<2.5e-7; Fisher exact test), consistent with a requirement for inactivation of the wildtype allele in the context of *BRCA1/BRCA2* deficiency^58^. An enrichment of signature 3 was also seen in *PALB2, FANCD2,* and *FANCM* germline PTV carriers (OR=3.7, *P*=0.01) including in pancreatic, ovarian, breast, and liver cancers. Interestingly, none of these cases showed evidence for somatic inactivation of the wildtype allele. In summary, these results suggest that mutagenesis attributable to signature 3 could be mediated by genetic defects in several members of the HR/ICL repair pathways.

**Figure 4.**
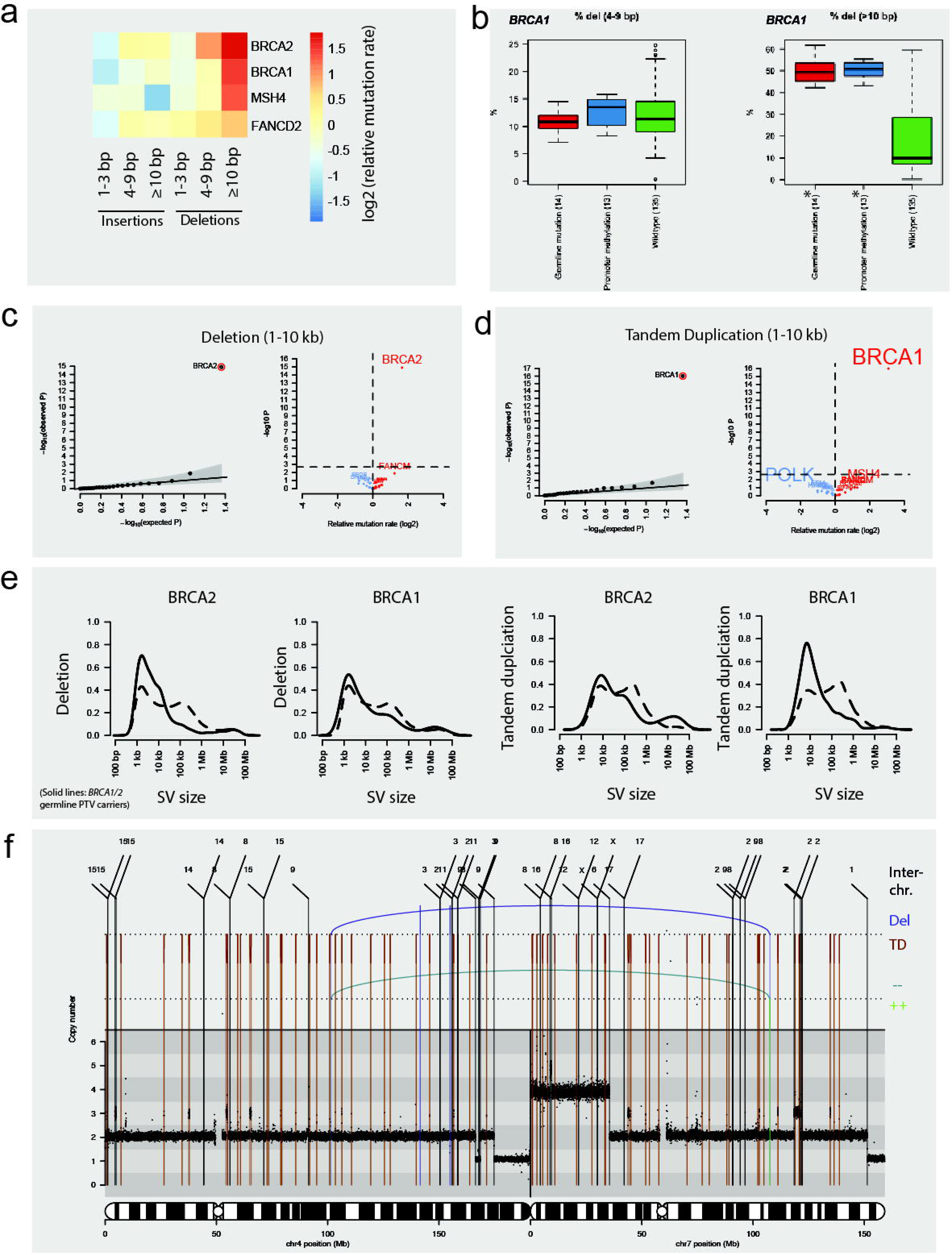
Association of germline PTVs with somatic indels and SVs. **(A)** Summary of germline PTV / somatic indel associations computed over all candidate genes represented by *N*≥4 germline PTV carriers in European donors, across PCAWG. (**B**) Association of epigenetic *BRCA1* gene silencing by promoter hypermethylation^65^ further substantiates specific somatic indel association with indel deletions >10bp (*denotes significance; 10% FDR). (**C**) QQ and volcano plot for SV deletions <10kb. (**D**) QQ and volcano plot for SV tandem duplications <10kb. (**E**) Somatic SV size spectrum shifts detected in association with germline PTVs in *BRCA1* and *BRCA2*, respectively (see also **Extended Data Fig. 7**). Dashed/solid lines: tumours lacking/harbouring germline PTVs in *BRCA1/2*. (**F**) Prostate cancer sample with numerous tandem duplications, which arose in a *BRCA1* germline PTV carrier. Abbreviations: Inter-chr., for inter-chromosomal; TD, for tandem duplication-type rearrangement; DEL, for deletion-type rearrangement; --/++, for inversion-type rearrangements (in the case of ++, reads point towards the q-telomere of a chromosome, and for -- towards the p-telomere, respectively).

Closer inspection of mutation spectra in tumours emerging in donors harbouring germline PTVs in these HR and ICL repair genes showed that, whilst there was evidence for an increased relative frequency of most mutation types in germline PTVs carriers, C>T substitutions at CpG dinucleotides were relatively depleted (**Fig. 2BCD**). Notably, analysis of absolute mutation counts revealed that C>T substitutions at CpGs occurred at a similar frequency in carriers of *BRCA1/2* germline PTVs as in donors lacking germline PTVs in *BRCA1/2* (**Extended Data Fig. 5**) – which suggests that the respective DNA lesions may not be repaired by these genes. C>T mutations at CpGs result from the spontaneous hydrolytic deamination of 5-methylcytosine, a widespread process represented by mutation signature 1 active in normal tissues and cancers^17,60^. Mutation signature 1 has been found in all cancer types studied to date^44^, and results in a relevant set of cancer driver mutations^61^, prompting us to more carefully follow up this finding.

**Figure 5.**
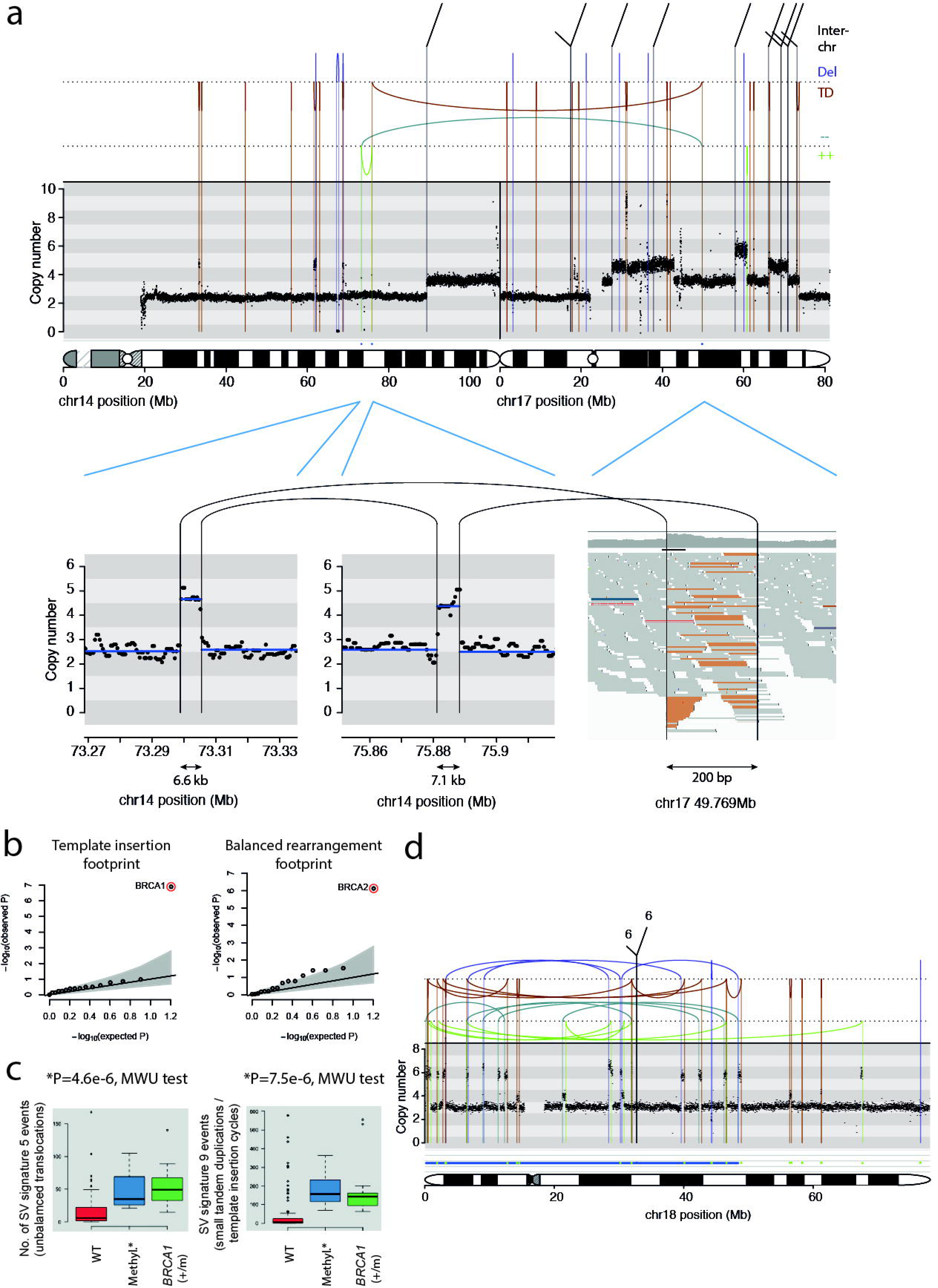
Association of germline PTVs with SV processes including complex patterns of SVs. **(A)** Example of a complex somatic SV in a breast cancer patient with *BRCA1* germline deficiency, with duplicated areas/templated insertions from distinct genomic regions. **(B)** QQ plot showing association of *BRCA1* germline PTVs with SVs exhibting cycles of templated insertions, and *BRCA2* germline PTVs with a balanced DNA rearrangement footprint. **(C)** Enrichment of an SV signature characterized by small tandem duplications and insertion cyles (“SV signature 9”)^66^ and an SV signature characterized by unbalanced translocations (“SV signature 5”) in *BRCA1* germline PTV carriers. Both observations are further corroborated by similar enrichments in ovarian and breast cancers showing *BRCA1* silencing via promoter hypermethylation (MWU for Mann-Whitney U test). (**D**) Highly complex rearrangement, potentially formed by DNA replication error-associated SV formation, in an ovarian cancer patient carrying a *BRCA1* germline PTV. Differing copy-number states suggest that duplications may have occurred on top of one another during SV formation.

### Germline PTV association with mutational signature 1: MBD4 germline PTV carriers show increased rates of a widespread clock-like mutational process

Intrigued by this observation, we performed a search for putative genetic determinants of the rate of mutation signature 1 – one of the most abundant mutational signatures across PCAWG tumor samples^45^ – based on germline PTVs within our set of 292 candidate genes. This analysis revealed *MDB4* as the only candidate gene associated with mutational signature 1 (relative rate (RR)=4; *P*<1e-5; **Fig. 3A**) implicating a gene that has not, to the best of our knowledge, previously been linked with the modulation of mutational processes in human cancer. We identified four carriers of diverse tumour types exhibiting heterozygous *MBD4* PTVs amongst European donors (**Table S3**). Separate quantification of C>T mutations at CpG sites in these carriers based on NpCpG motif analysis (**Supplemental Material**) further substantiated the association with signature 1 mutagenesis resulting in further increased significance (RR=5; *P*<1e-7; **Fig. 3B**). *MBD4* encodes a DNA glycosylase removing thymidines from T:G mismatches at methyl-CpG sites^62^, a functionality that may explain the germline-somatic association with mutation signature 1. The PTV-specific mutational type spectrum of *MBD4* demonstrated, accordingly, highly specific enrichment for all four Np[C>T]pG mutation types (**Fig. 3C**). Notably, analysis of 8,337 previously published TCGA WES samples^3,4^, which included 14 additional carriers of heterozygous germline *MBD4* PTVs, replicated the association of *MBD4* with somatic C>T transitions at CpGs (*P*=1e-3) based on nucleotide substitutions detectable in exome sequencing data (**Fig. 3D**).

Mutational signature 1 was recently shown to represent a ubiquitous clock-like mutational process the abundance of which is highly correlated with donor age in a variety of histologies^17^ – and for which, to the best of our knowledge, a germline genetic determinant has not previously been reported. To further analyse the nature of this genetic factor, we subsequently performed analyses of *MBD4* allelic state and gene expression, and investigated the genomic distribution of Np[C>T]pG substitutions in tumours emerging in *MBD4* germline PTV carriers. Two out of the four (50%) European donors harbouring *MBD4* germline PTVs displayed somatic loss of the *MBD4* wildtype allele (**Table S4**), which may have augmented signature 1 mutagenesis in the respective tumour specimens. Investigation of different genomic features including gene density, heterochromatin and DNA replication timing suggested that somatic Np[C>T]pG mutagenesis is augmented throughout the genome in association with *MBD4* germline PTVs (**Extended Data Fig. 6**).

**Figure 6.**
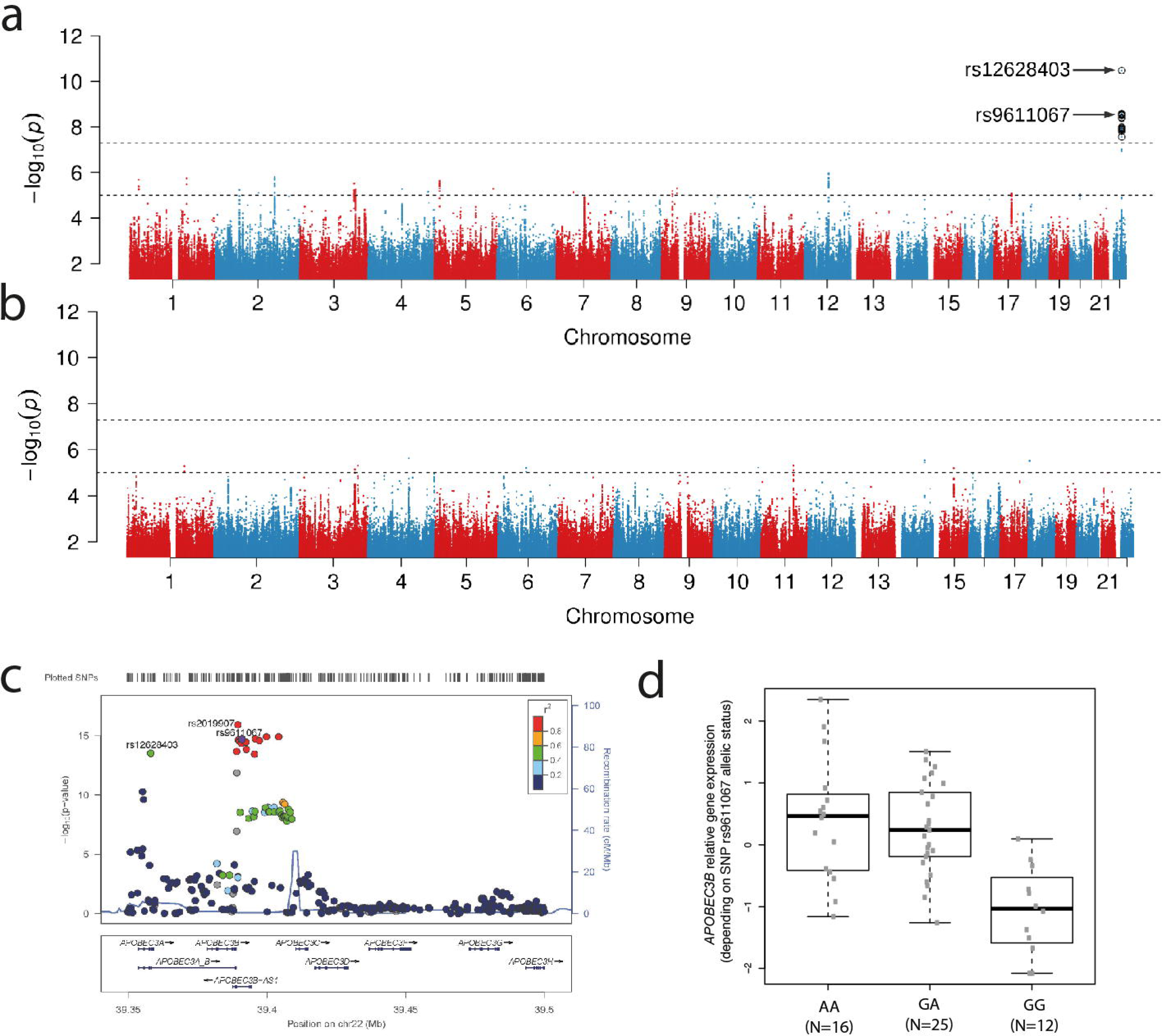
Association of common germline variants with APOBEC cytidine aminase attributable mutagenesis. **(A)** Manhattan plot for *APOBEC3B*-like signature enrichment in European PCAWG donors. **(B)** Manhattan plot for *APOBEC3A*-like signature enrichment in European PCAWG donors. **(C)** Close up of significant peaks, Manhattan plot for *APOBEC3B*-like signature enrichment in East Asian PCAWG donors showing two independent association signals at the APOBEC3 gene cluster. **(D)** *APOBEC3B* relative gene expression in liver cancer samples from Japan versus allelic status of rs9611067.

Gene expression profiling across cancers based on 1,220 PCAWG cancer samples with available transcriptome data^63^, notably, revealed an inverse correlation between *MBD4* gene expression levels in tumors and mutational signature 1 levels (*P*=1.9e-21; Mann-Whitney U-test; **Extended Data Fig. 6**), whereby tumours with low-level *MBD4* expression showed more signature 1 mutations. This result lends further support to the relationship between *MBD4* and signature 1, and suggests that *MBD4* expression differences may modulate levels of C>T mutations at CpGs across different tumour histologies.

### Germline PTV association with patterns of somatic indels and SVs

We next analysed indels, which can form at considerable rates in cancer^45,64^. Employing rare variant burden testing across cancers amongst the set of 292 candidate genes revealed associations between somatic indels and germline PTVs in *BRCA1*, *BRCA2*, *FANCD2*, and *MSH4* (**Fig. 4A**) – with *MSH4* clustering with the former three genes, all of which are involved in HR repair, also in terms of mutation type spectra (**Fig. 2E**). Whereas all four genes showed association with elevated levels of indels ≥10bp in size, only *BRCA2* associated with insertions and deletions 4-9bp in size, a finding further corroborated with ovarian and breast cancer samples exhibiting *BRCA1* promoter hypermethylation known to mediate *BRCA1* gene silencing^65^ (**Fig. 4B**, **Extended Data Fig. 7**). We additionally investigated germline PTVs in the context of microsatellite instability (MSI) (**Supplementary Notes**), identifying previously implicated genes of the DNA mismatch repair pathway^33^ as determinants for MSI (**Extended Data Fig. 8**).

**Figure 7.**
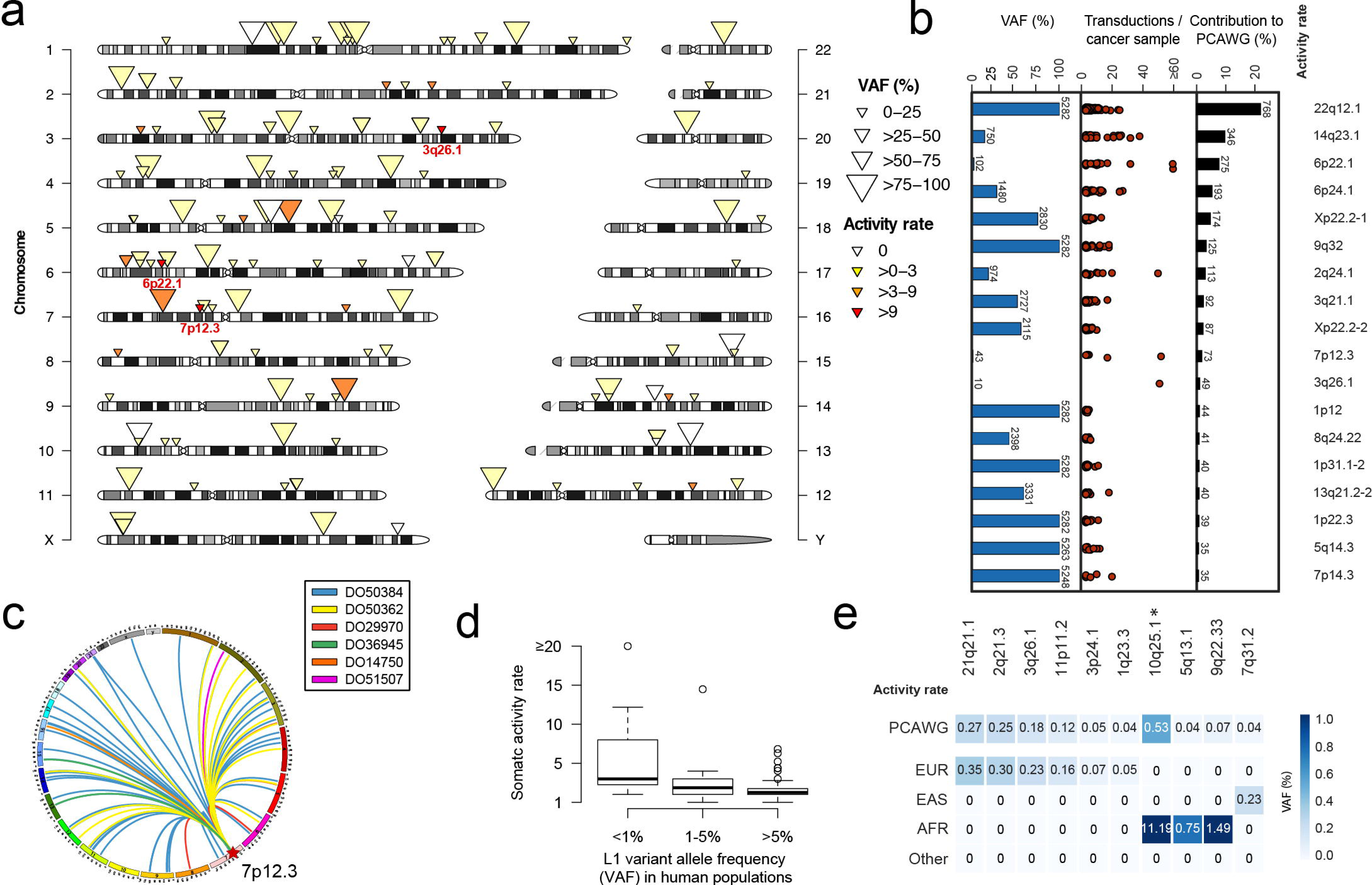
Germline L1 source elements driving somatic retrotransposition events in cancer. **(A)** Chromosomal map of germline L1 source elements with onferred somatic transduction activity. Elements with activity ‘0’ were identified as active in earlier studies, yet were not identified as active in PCAWG samples. (**B**) Source element contribution to L1-mediated transductions in PCAWG tumour samples versus variant allele frequency (VAF) in human populations (according to PCAWG germline genomic samples). The total number of transductions mediated by each source element and source element allele counts are shown as barplots. (**C**) A highly active germline L1 source element on chromosome 7 identified in donors of European and African ancestry (with VAF≤1%). (**D**) Higher inferred somatic L1 activities in rare compared to more common germline L1 source elements (*P*=0.0015; Kruskal-Wallis rank sum test). An outlying point observed for rare rare L1 source elements (VAF≤1%, inferred activity of 49) is shown at “≥20” (for visualization purposes). (**E**) L1 source elements seen in only one continental population. *Likely population-specific somatically active L1 source element present at appreciable frequency in African donors (VAF~11%) as well as in African 1000 Genomes Project samples (VAF~8%), yet entirely absent from European, East Asian and South Asian donors from PCAWG and the 1000 Genomes Project (see also Supplementary Note).

Somatic SVs, including balanced DNA rearrangements and copy-number alterations, represent the dominant form of mutation impacting cancer-related genes in several tumour types^6,45^. Employing rare variant association analysis across cancers, *BRCA2* germline PTVs associated with somatic deletion (**Fig. 4C**) and *BRCA1* germline PTVs with tandem duplication (**Fig. 4D**) rates, corroborating and further extending recent findings in breast cancer^18,42^. Deletion sizes were markedly reduced in *BRCA2* germline PTV carriers compared to sporadic tumours, whereas duplication sizes showed a marked reduction in *BRCA1* germline PTV carriers, an observation that we replicated in breast and ovarian cancers exhibiting *BRCA1* promoter hypermethylation (**Fig. 4E**, **Figure Extended Data Fig. 7**). Additionally, we detected a *BRCA1* germline PTV in a single prostate cancer patient whose tumour, notably, displayed >400 small tandem duplications (<100 kb) (**Fig. 4F**) ranking first among all prostate cancers in PCAWG (n=210) with regard to somatic tandem duplication load.

### Analysis of somatic SV processes implicates BRCA1 deficiency in DNA replication errors

Manual inspection of somatic SVs arising in *BRCA1* germline PTV carriers showed that these occasionally manifest as more complex rearrangements linking distant genomic regions including regions on different chromosomes (**Fig. 5A & Extended Data Fig. 9**). These SVs typically exhibited inserted fragments at their breakpoints, and closer inspection showed that these fragments corresponded to duplicates of existing genomic regions, suggesting that they became copied from a pre-existing template rather than resulting from concurrent DNA breaks that became fragmented and subsequently repaired (**Fig. 5A**). Using rare variant burden testing across cancers we searched amongst our set of 292 candidate genes for PTVs associating with SVs that exhibit a breakpoint footprint of templated insertions, which we identified using a novel computational approach^66^ (**Methods**). This analysis identified *BRCA1* as the only gene associated with a templated insertions footprint, whereas *BRCA2* germline PTVs showed association with a different footprint characteristic for balanced rearrangements (**Fig. 5B**).

The presence of templated insertions at SV breakpoints suggests the activity of a DNA polymerase and hence replicative processes during somatic SV formation in conjunction with *BRCA1* deficiency. Duplication of DNA sequence mediated by aberrant replication was previously studied primarily with respect to SV formation in congenital disorders^67-69^. In the context of aberrant DNA replication, template insertions have been attributed to low-processivity replication forks that switch templates and resume replication at a new site^68^. Complex rearrangements may arise in this context, given that relocated forks may themselves be prone to stalling and further template switching^67,68^.

To further corroborate our findings in conjunction with *BRCA1* deficiency, we addressed the question whether a particular ‘SV formation process’ generating complex DNA alterations may operate in *BRCA1*-deficient cancers. To do so, we made use of a computational approach evaluating the proximity of adjacent rearrangement junctions as well as of the orientation and order of joined segments^66^ to classify somatic SVs into signatures. Comparing *BRCA1* germline PTV carriers in ovarian cancer and breast cancer samples (representing 19 out of 27 *BRCA1* PTV carriers in PCAWG) versus sporadically arisen ovarian and breast cancers, we identified a preponderance of *BRCA1* PTV carriers to show two particular SV signatures. The first SV signature was characterized by relatively small tandem duplication events and cycles of templated insertions^66^ typically less than 100kb in size, implicating aberrant re-replication of genomic loci, and formation of SVs through template switching, in the context of *BRCA1* deficiency (**Fig. 5C**). The second SV signature was characterized by unbalanced translocations^66^, yet another form of somatic DNA rearrangement that appears to be increased in *BRCA1* deficient tumours. Reassuringly, the relationship of germline PTVs in *BRCA1* with these two signatures was further substantiated by a significant enrichment of the same two signatures in ovarian and breast cancers showing *BRCA1* promoter hypermethylation (**Fig. 5C**). We additionally analysed *BRCA2* germline PTV carriers using the same approach, identifying a preponderance of these to exhibiting three distinct somatic SV signatures^66^, including a signature characterized by deletions <10kb, a signature characterized by deletions of 10kb to 3Mb, and a signature characterized by reciprocal balanced rearrangements, respectively (*P*<0.05; Bonferroni correction).

Intriguingly, samples with genetic or epigenetic *BRCA1* inactivation also, occasionally, displayed much more complex rearrangement patterns – resulting in rearrangements that appear to involve series of highly inter-connected duplicative SVs. As one notable example, we detected 15 inter-linked relatively short duplicated segments in an ovarian cancer sample harbouring a germline *BRCA1* PTV (median segment size 75kb; minimum=21kb; maximum=232kb; **Fig. 5D**). While the marked complexity precludes classification with our computational approach^66^, there are intriguing parallels with the signature of short tandem duplications and templated insertion cycles seen enriched in a *BRCA1* deficient context. Both event types oscillate around few copy-number states and both appear to involve interlinked duplicative segments, whereby some of the duplications partaking in the complex rearrangement shown in **Fig. 5D** – presenting distinct duplicative copy-number states – may have occurred on top of one another. Based on these observations it is an intriguing possibility that a duplication-mediated process leading to massively complex rearrangements different from chromothripsis (chromosome shattering involving segment deletions/losses^22,24-27^ rather than segment duplications) may have formed this event. We searched for interconnected clusters of duplications involving at least ten relatively short (<500kb) interlinked duplicated segments (*i.e.* complex rearrangements resembling the event shown in **Fig. 5D**; **Supplementary Notes**). Nineteen such putative events were detected across all PCAWG donors, three of which arose in a *BRCA1*-deficient context, suggesting an enrichment of these complex rearrangement structures in *BRCA1*-defective tumours (OR=12.7; *P*=0.0028, Fisher’s exact test; **Fig. 5D, Extended Data Fig. 10, Extended Data File 1**). The interconnected rearrangements and the copy-number oscillating nature of these suggest that these structures likely formed in a single SV formation event, perhaps as an extreme outcome of erroneous DNA replication.

### Common haplotypes at 11q22 and 22q13 implicated in somatic mutagenesis across cancers

We next pursued a search for common germline polymorphisms (MAF>1%) associated with patterns of somatic mutation, by performing pan-cancer GWAS analyses for individual mutation types in European donors (**Methods**). These analyses identified a locus at 11q22 associated with overall mutation load at genome-wide significance (lead SNP rs12787749, MAF=1.1%, *P*=7.6e-11). This locus overlaps *YAP1*, an oncogene and effector of the Hippo tumor suppressor pathway^70^ (**Extended Data Fig. 11**).

We next considered mutational signatures, focusing on determinants of mutations attributed to the activity of the APOBEC family of cytidine deaminases, for which genetic determinants had previously been detected via locus-targeted analysis^32,71^. GWAS analysis showed no pan-cancer association for mutational signatures (2 and 13) attributed to APOBEC activity^44^ in Europeans (**Extended Data Fig. 11**). Both signatures 2 and 13, however, represent composites of the enzymatic activity of *APOBEC3A* and *APOBEC3B*, which mediate mutagenesis in cancer at different activity levels^72^. We therefore refined this analysis by decomposing APOBEC mutation signatures into *APOBEC3A*-like (YT<u>C</u>A) and *APOBEC3B-*like (RT<u>C</u>A) signatures^72^, computing signature enrichments in a tetranucleotide context^72^ (**Supplementary Notes**). Using the subset of PCAWG samples (*N*=1300) showing *APOBEC3A*-like and *APOBEC3B*-like signature enrichments (**Supplement**), we identified common genetic variants that associated with *APOBEC3B*-like signature enrichments at genome-wide significance (**Fig. 6AB**). The most strongly associated SNP was rs12628403 (*P*=3.35×10^−11^), which tags a deletion polymorphism fusing *APOBEC3A* with noncoding regions of *APOBEC3B* – a variant previously implicated in APOBEC mutagenesis and breast cancer risk^32,71^. Additionally, we identified a second locus, downstream of *APOBEC3* (lead SNP rs9611067, ~300kb from rs12628403, *P*=2.89×10^−9^). We also observed strengthened associations amongst donors of East Asian ancestry (*N*=379), in whom the *APOBEC3A/B* deletion polymorphism is more frequent (*P*=2.82×10^−14^ for rs12628403, *P*=1.87×10^−15^ for rs9611067; **Fig. 6C**, **Extended Data Fig. 11**).

Notably, *cis*-expression quantitative trait locus (eQTL) mapping pursued in lymphoblastoid cell lines^73^ previously associated rs9611067 with *APOBEC3B* and *APOBEC3A* expression. Pan-cancer transcriptome analysis, which we performed across 1,047 PCAWG tumour samples with transcriptome data, showed that this eQTL extends to tumours with highest effect sizes measured for *APOBEC3B* (**Fig. 6D, Extended Data Fig. 12**). rs9611067 and rs12628403 are in strong linkage disequilibrium (LD) in Asians but not Europeans (*r^2^*=0.07 in Europeans and *r*^2^=0.41 in East Asians); analyses controlling for rs12628403 weakened but did not eliminate the association for rs9611067 (*P*=4.7e-05; see **Supplement**). Closer inspection of the relationship of these SNPs with *APOBEC3B-*like signature enrichment, notably, showed a more pronounced genetic effect for relatively low-to moderate-level than for high-level *APOBEC3B*-like signature enrichment, perhaps since samples with higher *APOBEC3B-*like mutagenesis are more prone to be influenced by environmental factors that mask genetic factors (**Extended Data Fig. 12**). Interestingly, we observed a reduction of viral (hepatitis B or C) infections amongst cancer samples from the Japanese liver cancer cohort, the largest tumor-type specific cohort in PCAWG^74^, in carriers of the rs9611067 risk allele (OR=0.29; *P*=0.02, Fisher’s exact test; **Table S5**). These findings suggest a possible interaction between APOBEC mutagenesis and infection (a potentially relevant environmental factor), or may result from APOBEC mutagenesis and viral infection both independently contributing to the disease.

### Over one hundred germline L1 source elements associated with somatic retrotransposition in cancer

L1/LINE elements are a germline SV class with the ability to result in widespread somatic mutation through retrotransposition^19,21,75,76^, representing the third most abundant class of somatic SV in cancer^45^. L1 elements are a resource for cancer-driving mutations, which can involve somatic insertion-associated Megabase-sized DNA rearrangements and other mechanisms^77-79^. L1 retrotransposition in cancer occurs through the activity of an as yet largely undetermined set of L1 germline source loci, which we here aimed to characterize leveraging the unprecedented set of 2,642 paired tumour and normal whole genomes. To trace somatic L1 events to individual germline source loci we searched for small tracks of L1-adjacent unique (non-repetitive) DNA sequences mobilised through L1 retrotransposition^19,80-82^, known as somatic L1-mediated transductions^19,80-82^, across all cancer samples in PCAWG.

Doing so, we inferred the somatic activity of 124 germline L1 elements, including 75 that represent insertions with respect to the human reference genome, as well as 49 elements that are part of the reference assembly (**Fig. 7A**, **Table S6**). The majority (55%; 41/75) of polymorphic germline L1 source elements identified as insertions relative to the reference were novel with respect to current databases (**Table S6**). Irrespective of continental population ancestry, PCAWG donors typically exhibited 50–65 germline source elements with the ability to promote retrotransposition in cancer (**Extended Data Fig. 13**). We identified SNPs in strong LD (defined^47^ as *r*^2^≥0.8) for the majority of somatically active source elements inferred as polymorphic insertions (*i.e.*, 53/68 germline L1s with MAF≥0.1%), which will facilitate identifying these active L1 source loci in future cancer genomics studies (**Table S7**).

Overall, we inferred 2,923 L1-mediated transductions amongst 20,230 somatic retrotranspositions events identified among PCAWG donors^77^. 43% of cancer samples showed evidence for somatic retrotransposition, with notable enrichments for tumour types with high source element activity such as lung squamous carcinoma, oesophageal adenocarcinoma, and colorectal adenocarcinoma^77^. We observed notable differences in activity amongst L1 source elements. Sixteen moderate to high activity source loci contributed at least three transduction events on average in cancer genomes with detectable L1 activity, including rare L1 germline polymorphisms such as a novel element on chromosome 7p12.3 (variant allele frequency (VAF)≤1% in European and African populations) giving rise to 75 transductions amongst only six cancer samples with L1 activity (**Fig. 7BC**). By comparison, a previously described 22q12.1 source element^19,83^ detected across all donors of European, East Asian, African, South Asian and Native American ancestry exhibited more modest activity – yet owing to its fixed status contributed 22% of the transductions in PCAWG overall (**Fig. 7B, Extended Data Fig. 13**). Curiously, more active L1 source elements showed a preponderance towards rarer human population frequency (*P*=0.0015; Kruskal-Wallis rank sum test), with the rarest L1 source elements (VAF<1% across human populations) displaying ~3.5-fold higher activity than the most common source elements (VAF>5%) (**Fig. 7D**). According to classical theories for transposon evolution^84^, differences in the duration and intensity of selective pressures acting upon rare versus more common L1 elements may serve to explain this observation. Notably, some L1 germline source loci demonstrated signs of population-specificity, including a somatically active L1 source element on chromosome 10q25.1 with VAF=11% in donors of African ancestry, yet entirely absent in PCAWG donors of European and Asian ancestry as well as in 1000 Genomes Project^40,50^ donors of European and Asian ancestry (**Fig. 7E** and **Supplementary Notes**).

## Conclusions

We report a novel resource comprising 88 million germline variants in 2,642 widely available cancer genomes, and demonstrate patterns of somatic mutation associated with germline PTVs, common SNPs and L1 source elements across diverse cancer types, which includes an association seen for a presumably population-specific somatically active L1 source element occurring in donors of African ancestry.

Findings from our study reveal common genetic variants associated with somatic base substitution rates and processes at genome-wide significance, namely those associated with mutagenesis attributable to *APOBEC3B*. Additionally, we implicate *BRCA1* germline PTV in complex SV patterns characterized by duplication-associated rearrangements likely mediated by aberrant DNA replication, an SV formation process that has remained under-studied in cancer. Replication-associated somatic SV formation may thus be more common than currently thought, and the *BRCA1* gene may, in its intact form, serve to protect cells from such mutational events. Additionally, complex patterns of alterations we identified in conjunction with *BRCA1*-defects may facilitate accelerated karyotype evolution. By comparison, we implicated *BRCA2* germline PTVs with somatic SV formation events likely to be initiated by DNA double strand breaks, including deletions and balanced DNA rearrangements.

Genes implicated in augmenting somatic mutation processes include some not previously reported as cancer predisposition genes. For example, our study implicates *MSH*4 with indel patterns and somatic base substitution spectra in cancer genomes similar to those seen in conjunction with HR repair deficiency (*i.e. BRCA1* and *BRCA2* PTVs), an observation that warrants follow-up studies given the relevance of HR repair deficiency for therapeutic targeting^41^. Evidence that *MSH4* can augment the formation of short deletions, notably, has also been presented in yeast recently^85^. Elevation of mutation signature 1 in association with *MBD4* germline PTV is, to our knowledge, the first demonstration of a heritable factor mediating inter-individual variation in the spontaneous hydrolytic deamination of cytosines at CpGs (which are commonly methylated) in humans. Signature 1 shows clock-like properties in tumour and normal tissues^8,17,60,86^ and may signify the number of post-zygotic divisions a cell has undergone^17^. Given its abundance across tissues^8,17,60,86^ and its correlation with patient age^17^, it has been suggested that signature 1 may influence ageing-related illnesses such as cancer^8^. If that is the case, this would render *MBD4* a novel candidate risk gene for age-related pathologies including (and not limited to) cancer. Notably, *MBD4* knockout (*Mbd4−/−*) in mice was found to promote gastrointestinal tumour formation, and mutation analyses pursued by targeting a ~500bp-sized locus are consistent with increased spontaneous C>T transitions at CpGs in *Mbd4−/−* mice^87^, further supporting our findings based on human cancer genomes. Signature 1 partially recapitulates the pattern of *de novo* mutations in the germline^17^, raising the question whether *MBD4* germline PTV carriers may show increased germline mutations. Efforts to systematically catalogue *de novo* mutations in families^88,89^ could address this question in the future.

Our study also has remaining limitations. Germline-somatic associations uncovered in our study mainly pertain to mutational patterns seen in a large fraction of donors, and depend on the number of germline variant carriers sampled in individual tumour types – a number that in spite of the inclusion of >2600 whole cancer genomes in our resource is inevitably going to be too small for candidate genes in which germline PTVs are relatively rare. In line with this, controlling the FDR at 20% (at “near significance”) let to a number of additional credible ‘hits’, including *PALB2* and *RECQL*, which notably exhibited elevated indel and base substitution spectra similar to other HR genes (**Extended Data Fig. 14, Fig. 2DE**). In the future, increased samples sizes are likely to provide insights into currently unexplained mutational signatures, including such restricted to particular tissues, on the basis of the approaches devised in our study. Mutation in cancer reflects the cumulative activity of different mutational processes, and our study identified a variety of associations between germline variants and somatic mutation, highlighting how variation in the germline can markedly shape cancer genomes.

Finally, our dataset will likely show utility as a reference for prioritisation of rare pathogenic variants and imputation of common variants in cancer-focused disease studies, which is why we are releasing the full dataset (including genotype likelihoods) as a resource to the cancer community through standard data repositories (see “Data availability” below). The availability of both germline and somatic variants for each donor will facilitate utilisation of this reference resource for research on somatic mutation processes and putative germline cancer susceptibility variants.

## Data availability

All data of our germline variant resource, including sequence alignment (BAM) files and haplotype-block phased variant call files (VCF), are made available to the community via ICGC and TCGA-associated PCAWG data portals^45^ (http://docs.icgc.org/pcawg) using controlled data access principles that have become standard in the cancer genomics community. Data releases can be accessed from the TCGA (dbGAP accession phs000178), the EGA (EGAS00001001692), and the ICGC (https://dcc.icgc.org/releases/PCAWG).

## Acknowledgements

This project received funding by the European Commission (EC) through an ERC Starting Grant to JOK, the EurocanPlatform project of the EC (to JOK and AB), the European Open Science Cloud (EOSC) pilot project of the EC (to JOK and SN), the European Union’s H2020 research and innovation programme under grant agreement No 635290 (PanCanRisk to XE, SO and OS) and Catalan Research Groups (AGAUR) Generalitat de Catalunya, projects funded by the German Federal Ministry of Education and Research (the Biotop project, to JOK; the ICGC DE mining project, to JOK and RS; de.NBI project, to JOK), the Spanish Ministry of Economy and Competitiveness, ‘Centro de Excelencia Severo Ochoa 2013-2017 and the CERCA Programme / Generalitat de Catalunya. S.M.W. received funding through a SNSF Early Postdoc Mobility fellowship (P2ELP3_155365) and an EMBO Long-Term Fellowship (ALTF 755-2014). We thank Po-Ru Loh (Harvard T.H. Chan School of Public Health) for support in the use of EAGLE2 for phasing. We further thank the following providers/facilities for principally enabling and supporting the processing >2600 cancer whole genomes: Annai Systems – Annai Systems BioCompute Farm; Broad Institute – Broad Institute’s compute clusters; DNAnexus – DNAnexus Cloud Platform (kindly acknowledging Andrew Carroll); EMBL – Embassy Cloud, Heidelberg IT and GeneCore compute clusters. We further acknowledge the teams of Seven Bridges Genomics Inc., Fujitsu Inc., Intel Inc., and SAP Inc. for their invaluable support in cloud and high-performance computing. The results reported here are mainly based upon data generated through the ICGC (www.icgc.org) and TCGA (http://cancergenome.nih.gov) and we acknowledge the specimen donors as well as the various research groups involved in sampling and sequencing.

## Author contributions

Germline SNP/Indel callsets: GT, SW, SS, AS, ADB, PS, EG, TS, YW, MW, SY, JK, FV; HLA genotyping: SH, SI, HN; Merging and phasing of germline variant release set: TR, OD, SW, DW, JK; Validation of SNP/Indel calls: FM, SW, LD, JS, SO; Microarray based verification of germline SVs: TR; Germline SV callset: TR, SY, GD, JW, SO, JK; Germline MEI callset and L1 analysis: BRM, JT; Analysis of cancer risk genes: SW, RS, DE; Germline-somatic association analyses for rare variants: SW, EP, DE, JK; Germline-somatic association analyses for common variants: BZ, LS, XH, SW, OS, JK, SC, NC; Germline risk association analyses: RR, GE, HS, SO, JC, AB, OH, XE; Interpretation of mutational processes involving base substitutions: SW, DG, SR, NS, JK; Interpretation of indel processes: SW, EP; Interpretation of SV processes: SW, YL, TR, JK; Cloud workflow management and shepherding: SY, IL, AC, SN; Ancestry analysis: SS, MW, YW, SW, GT, CB, CDB, FV; Coordination of PCAWG germline working group: NH; Project supervision: FV, OH, JT, BZ, DE, XE, JK; The paper was written by SW, DE, XE and JK, with individual contributions and input from the other authors; PCAWG germline working group co-chairs: XE, JK. All authors contributed to data interpretation.

